# NvERTx: A gene expression database to compare Embryogenesis and Regeneration in the sea anemone *Nematostella vectensis*

**DOI:** 10.1101/242370

**Authors:** Jacob F. Warner, Vincent Guerlais, Aldine R. Amiel, Hereroa Johnston, Karine Nedoncelle, Eric Röttinger

## Abstract

For more than a century researchers have been comparing embryogenesis and regeneration hoping that lessons learned from embryonic development will unlock hidden regenerative potential. This problem has historically been a difficult one to investigate since the best regenerative model systems are poor embryonic models and vice versa. Recently however, the comparison of embryogenesis and regeneration has seen renewed interest as emerging models including the sea anemone *Nematostella vectensis* have allowed researchers to investigate these processes in the same organism. This interest has been further fueled by the advent of high-throughput transcriptomic analyses that provide virtual mountains of data. Unfortunately much of this data remains in raw unanalyzed formats that are difficult to access or browse. Here we present ***N****ematostella* ***v****ectensis* **E**mbryogenesis and **R**egeneration **T**ranscriptomi**cs** - NvERTx, the first platform for comparing gene expression during embryogenesis and regeneration. NvERTx is comprised of close to 50 RNAseq datasets spanning embryogenesis and regeneration in *Nematostella*. These data were used to perform a robust *de novo* transcriptome assembly which users can search, BLAST and plot expression of multiple genes during these two developmental processes. The site is also home to the results of gene clustering analyses, to further mine the data and identify groups of co-expressed genes. The site can be accessed at http://nvertx.kahikai.org.

## Introduction

A long-standing question in the field of regeneration is to what extent regenerative programs recapitulate development. Comparing gene expression during these two processes provides clues as to how genes activated during embryogenesis are re-deployed during regeneration. The majority of these studies are limited to individual or groups of genes that are assayed in specific regenerative contexts (Binari et al., 2013; Carlson et al., 2001; Gardiner et al., 1995; Imokawa and Yoshizato, 1997; Kaloulis et al., 2004; Katz et al., 2015; Özpolat et al., 2012; Reitzel et al., 2007; Torok et al., 1998; Wang and Beck, 2014). To date, no study has systematically compared embryogenesis to whole body regeneration on a global transcriptomic level. One organism however, is especially amenable to this line of study: the sea anemone *Nematostella vectensis* (Fig. 1A) (Layden et al., 2016; Reitzel et al., 2007).

**Figure 1:**
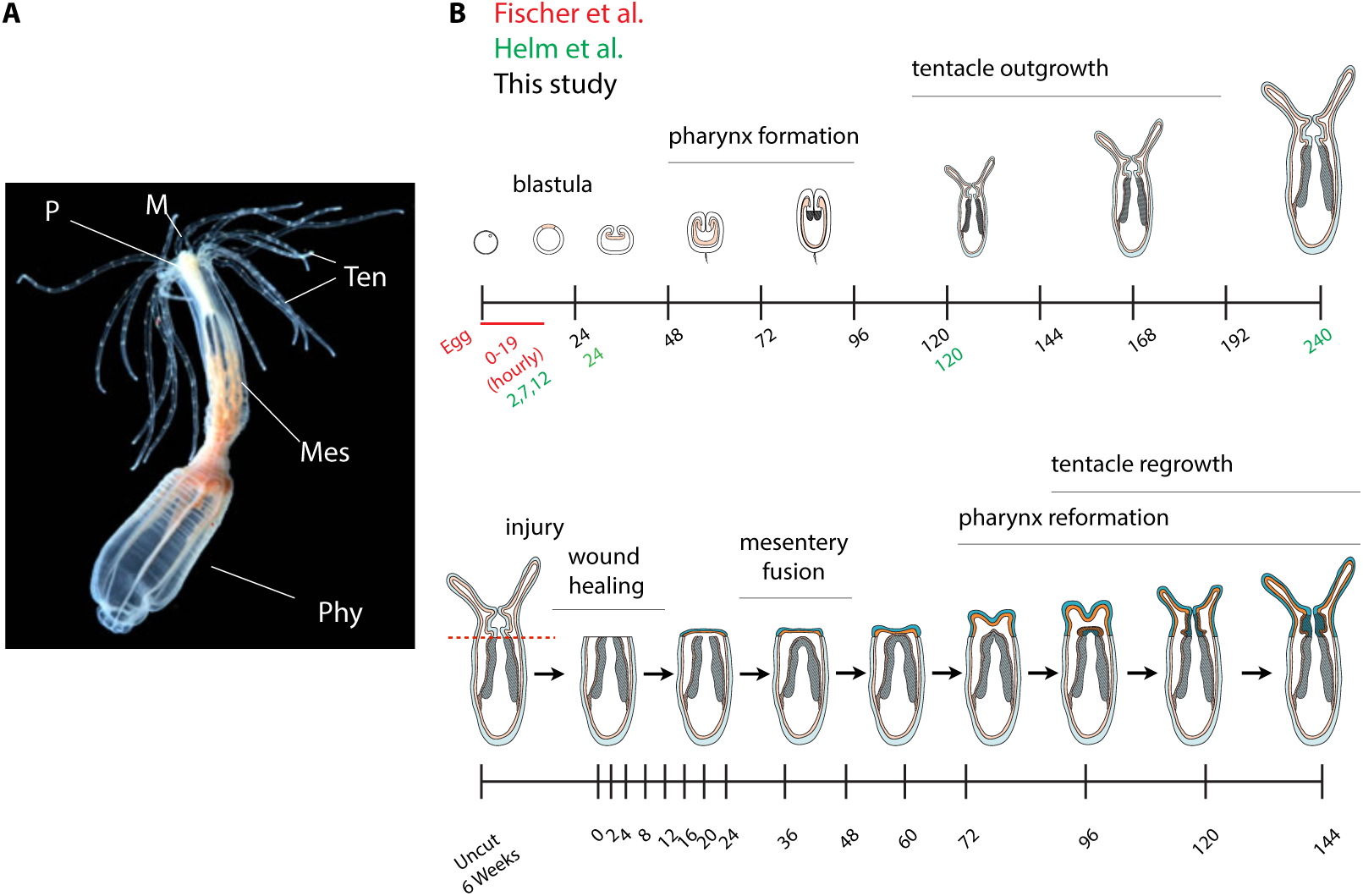
A) *Nematostella* anatomy. *Nematostella* is a small sea anemone (~5cm) comprised of a mouth (M), pharynx (P) tentacles (Ten) and a body column with internal structures called mesenteries (Mes), the posterior section of which is termed the physa (Phy). B) Schematic of RNAseq samples included in the NvERTx database. Three datasets spanning embryogenesis are included: Fischer et al. (2014) sampled hourly from 0-19hpf, Helm et al. (2013) sampled 2, 7, 12, 24, 120, and 240hpf, and this study sampled daily from 24 to 192hpf. Regeneration was sampled from six-week-old animals at uncut, 0, 2, 4, 8, 12, 16, 20, 24, 36, 48, 60, 72, 96, 120, and 144 hpa.

*Nematostella* has long been used as a model system for embryonic development. *Nematostella* reproduce sexually and after fertilization the zygote undergoes a series of cleavages to form a blastula. Gastrulation occurs at the animal pole of and shortly thereafter the embryo enters a swimming planula stage during which the pharynx and internal structures, termed mesenteries, develop. After several days, this planula settles, develops tentacles, and enters a juvenile stage (reviewed in: Layden et al., 2016). *Nematostella* development research entered the age of genomics with the sequencing of its genome by Putnam and colleagues in 2007 (Putnam et al., 2007; reviewed in: Technau and Schwaiger, 2015). Since then, a large number of developmental genes have been identified in *Nematostella* and many commonalities between *Nematostella* and bilatarian development have emerged (Amiel et al., 2017; Genikhovich et al., 2015; Layden and Martindale, 2014; Layden et al., 2012; Leclère et al., 2016; Matus et al., 2008; Meyer et al., 2011; Röttinger et al., 2012). With the advent of high-throughput transcriptomics, several studies have examined gene expression during embryogenesis at the whole-genome level (Helm et al., 2013; Tulin et al., 2013) firmly establishing *Nematostella* as an embryonic model.

More recently, *Nematostella* has shown itself to be a powerful model for regeneration. Upon bisection, *Nematostella* is capable of regenerating the missing body half after approximately 6 days post amputation (Bossert et al., 2013). Following sub-pharyngeal amputation (head removal) regeneration occurs via a highly dynamic process: First there is an initial wound healing phase of approximately 6 hours, then regeneration follows a stereotypic program in which the mesenteries fuse and via subsequent cell-proliferation reform the missing pharynx and tentacles over the course of 6 days (Amiel et al., 2015; DuBuc et al., 2014). This process is known to use several developmental signaling pathways originally deployed during embryogenesis (DuBuc et al., 2014; Passamaneck et al., 2012; Schaffer et al., 2016; Trevino et al., 2011). It remains unclear, however, if these pathways are deployed the same way, i.e. with similar or divergent regulatory logic. One way to address this question is to systematically compare gene expression profiles during embryonic development and regeneration to identify groups of genes originally used during embryogenesis that are re-used during regeneration. To facilitate these studies we created ***N**ematostella* **v***ectensis* **E**mbryogenesis and **R**egeneration **T**ranscriptomi**cs** - NvERTx: A quantitative gene expression database for comparing embryogenesis and regeneration in the sea anemone *Nematostella vectensis*. NvERTx is comprised of several datasets spanning embryogenesis (Helm et al., 2013; Tulin et al., 2013)(this study) and regeneration (this study). We used pooled RNAseq data from *Nematostella* embryogensis and regeneration to generate a *de novo* transcriptome assembly. Using this assembly, we then quantified each of the RNAseq datasets. We then clustered the transcripts to discover groups of genes that share similar expression. All of these data can be found in a searchable database that is accessible at www.nvertx.kahikai.org. This tool can be used to find transcript sequences, identify co-expressed genes, and directly compare expression profiles during embryogenesis and regeneration.

## Results and Discussion

### NvERTx Database: de novo assembly, annotation, and data quantification

NvERTx is a quantitative gene expression database for embryogenesis and regeneration (Fig. 1C) comprised of several RNAseq datasets. It includes previously published datasets spanning very early embryogenesis to polyp from Helm et al. (sampled 2, 7, 12, 24, 120 and 240 hours post fertilization (hpf))(Helm et al., 2013) and Fisher et al. (sampled 0, 1, 2, 3, 4, 5, 6, 7, 8, 9, 10, 11, 12, 13, 14, 15, 16, 17, 18, and 19 hpf) (Fischer et al., 2014). To complement these, and sample timepoints during tentaclegenesis and pharynx formation, we generated an additional dataset sampled at 24, 48, 72, 96, 120, 144, 168, 192 hpf. Together these datasets cover the major hallmarks of *Nematostella* development including blastula stages (12-24 hpf), gastrula stages (24-48hpf), planula stages (48-120hpf) and juvenile stages (120-244hpf). The regeneration RNAseq data were sampled from six week old juveniles after sub-pharyngeal amputation at uncut, 0, 2, 4, 8, 12, 16, 20, 24, 36, 48, 60, 72, 96, 120 and 144 hours post amputation (hpa) (see Fig. 1B for sampling strategy). We chose these timepoints because they span the most important events of regeneration including wound healing (0-6hpa), pharynx reformation (24-72hpa) and tentacle reformation (72-120hpa)(see Fig. 1A for Nematostella anatomy)(Amiel et al., 2015), all stages for which we have embryonic data spanning the initial development of these structures. For each of these datasets, we obtained the raw sequencing reads and used these as input into our quantitative workflow.

To quantify the RNAseq data, we performed a *de novo* transcriptome assembly, which we term NvERTx. We assembled this transcriptome using the short read assembler Trinity with combined paired-end reads from our regeneration dataset and additional embryonic data from Tulin et al. sampled at 0, 6, 12, 18, 24 hpf (Haas et al., 2013; Tulin et al., 2013) as input (see materials and methods for complete workflow). In the resulting assembly we identified 234381 transcripts and designated each a unique NvERTx.4 number. We then annotated this transcriptome by extracting the predicted coding sequences using OrfPredictor (Min et al., 2005) and comparing the resultant protein sequences to NCBI’s non-redundant protein database (nr) and the Uniprot database using the blast-like tool PLAST (Nguyen and Lavenier, 2009). From these analyses we identified 85475 transcripts with a significant hit to nr (e value < 5e-5) and 69335 transcripts with a significant hit to the Uniprot databases (e value < 5e-5). Additionally we compared each transcript to the current Nemve1 gene predictions (https://genome.jgi.doe.gov/Nemve1/Nemve1.home.html) using BLASTn and identified 102581 transcripts with a significant hit (e value < 5e-5) (Kent, 2002). For each of the 234381 NvERTx transcripts we provide all available annotations. We then used the transcriptome assembly to quantify each of the RNAseq datasets. We did this by aligning the reads using Bowtie2 (Langmead et al., 2009) and quantifying each transcript using RSEM and edgeR(Li and Dewey, 2011; Robinson et al., 2010). Transcripts with the same best Nemve1 hit were combined to obtain ‘gene-level’ quantification. Finally, the three embryonic datasets were corrected for batch effects using the sva R package with time-point as a categorical covariate (Leek et al., 2012)(see materials and methods for details). The quantified datasets are reported for each transcript in the database.

### Retrieving expression plots, count tables and sequences

The NvERTx database can be accessed using multiple points of entry: by searching the annotations (Fig. 2Ai), using the BLAST tool (Fig.2Bi) or exploring the co-expression clusters (Fig. 2Ci)(see section: *Exploring Gene Expression Clusters*.) To demonstrate this, we use the *Nematostella* Brachyury protein (*Nvbra*, genebank ID: AAO27886.2) as an example. To retrieve transcripts corresponding to *Nvbra* we can use the search function to query the transcript annotations (Fig. 2Ai). To search for *Nvbra*, we can use the Nemve1 gene model ID from the current genome assembly (jgi|Nemve1|770), the NCBI GenPept accession number (gi| 122058623), the NCBI GenBank accession number (AAO27886.2), or the gene name (‘Brachyury’) (Fig. 2Ai). Using any of these queries identifies several transcripts that correspond to *brachyury* (Fig. 2B). Multiple matching transcripts reflect the different isoforms predicted by the transcriptome assembler. We can confirm which transcripts correspond to *Nvbra* by examining the annotations that are reported with the search results. It is normal for different isoforms to have slightly different annotations as each transcript was individually annotated. Clicking the NvERTx.4 numbers fills in the field on the left of the screen, enabling the user to directly compare their expression profiles (Fig. 3A).

**Figure 2:**
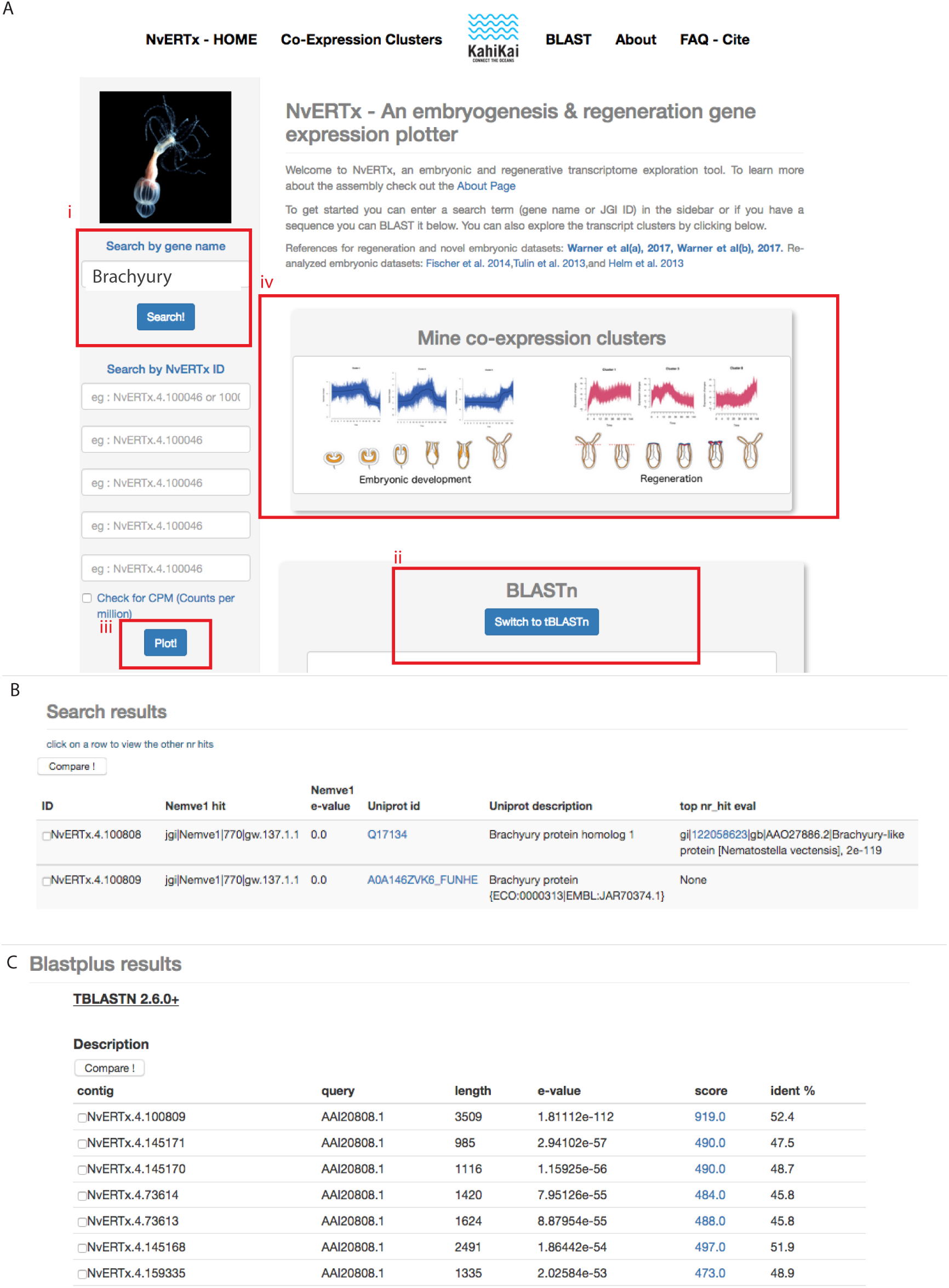
A) Screenshot of the NvERTx portal. **Ai**: Users can search for genes using the gene name, Nemve1 accession number, or NCBI genebank accession number. **Aii**: The transcriptome can also be searched using BLASTn or tBLASTn. **Aiii**: Multiple transcripts can be queried simultaneously. **Aiv**: Users can also directly explore co-expression clusters from embryogenesis and regeneration to identify groups of co-expressed genes. B) Screen shot of results from searching the annotations using the term “Brachyury”. C) tBLASTn results using *Mus musculus* Brachyury as a query (genbank AAI20808.1) identifies several homologous transcripts. The top scoring isoform is reported first.

**Figure 3:**
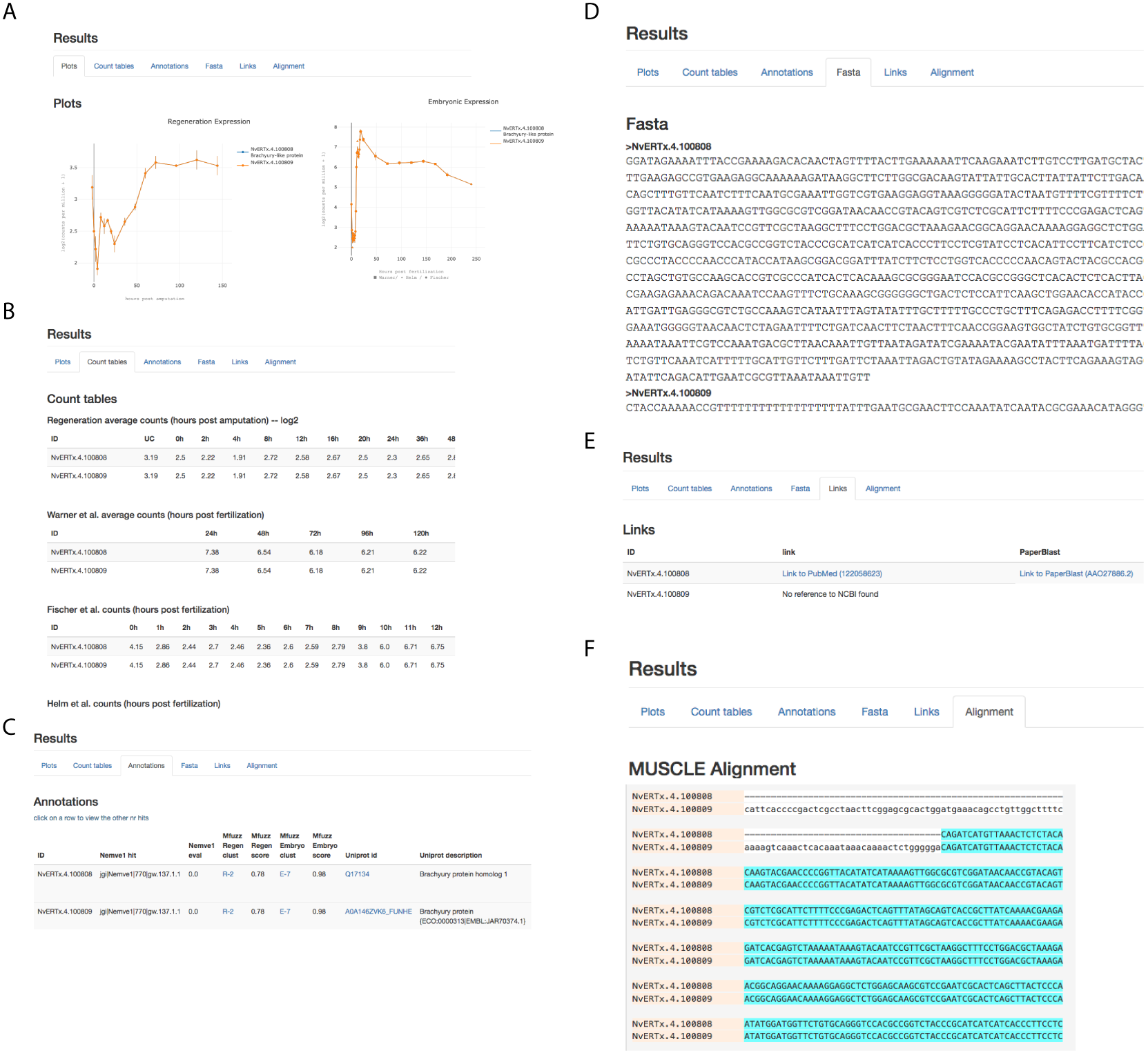
Example results for the transcripts NvERTx.4.100808 and NvERTx.4.100809. A) Expression plots for regeneration (left) and embryogenesis (right). The two transcripts are isoforms of the same genes so their expression profile plots and counts are equivalent. B) Count data from each of the datasets. C) Transcript annotations. D) Sequences in FASTA format. E) Bibliographical resources including Pubmed links and PaperBlast queries. F) MUSCLE alignment to compare similar transcripts. The alignment shown is cropped to display the homologous region.

Similarly the BLAST tool can be used to retrieve NvERTx.4 transcripts using either a nucleotide (BLASTn) or protein (tBLASTn) query. The reported alignments can be then used to identify the correct transcripts. For example using tBLASTn against the transcriptome with *Mus Musculus* Brachyury (NCBI GenBank accession number AAI20808.1) we find several homologous transcripts, the first of which is *Nvbra* (Fig. 2C). Again clicking the NvERTx.4 number fills in the field on the left of the screen enabling the user to obtain the expression plots and annotations.

Once the NvERTx.4 IDs for *Nvbra*, NvERTx.4.100808 and NvERTx.4.100809, are selected we can query the database by clicking ‘Plot!’ on the left (Fig. 2Aiii). The first page that appears displays the transcripts expression during regeneration and embryogenesis (Fig. 3A). We can see that the expression of *Nvbra* exhibits two peaks during regeneration beginning at 8hpa and 60hpa while during embryogenesis *Nvbra* is expressed early, rapidly peaks at 20hpf, then decreases throughout development. Note that the expression profiles for the two transcripts are perfectly super-imposed and appear as one. This is because transcripts corresponding to the same Nemve1 best hit are quantified equivalently as they are from the same ‘gene’. To distinguish transcript isoforms from separate genes we can compare the individual transcripts in the ‘alignment’ tab where a MUSCLE sequence alignment is reported (Edgar, 2004)(Fig. 3F). In this case we observe that NvERTx.4.100809 is a longer assembled isoform of *Nvbra*. The results tabset also includes the normalized count tables (Fig. 3B), transcript annotations (Fig. 3C), the sequences in FASTA format (Fig. 3D), and links to bibliographical resources including PubMed articles citing the protein and a PaperBlast query (Price and Arkin, 2017)(Fig. 3E). In the annotations tab, we can also see which co-expression cluster the transcript belongs to for embryogenesis and regeneration. Exploring these clusters is very useful for identifying co-expressed genes.

### Exploring Gene Expression Clusters

Co-expression analysis is a particularly useful way to identify genes that function in the same gene regulatory module. Co-expressed genes can also represent groups of genes that participate in a similar biological function. One method to identify co-expressed genes is to cluster genes or transcripts by expression profile. For NvERTx, we performed fuzzy c-means clustering to regroup genes by expression profile and provide those clusters in the ‘Co-Expression Clusters’ section of the site. These clusters can be used to identify transcripts that are co-expressed with a given gene of interest. Furthermore comparing the membership of gene-clusters during embryogenesis and regeneration can be used to identify groups of genes that function similarly during these processes.

The gene expression clusters can be browsed by either clicking a cluster in the ‘Co-Expression Clusters’ section (Fig. 5) or by following a direct link from the annotation table of a transcript to its cluster (Fig. 3C). Using our previous example, *Nvbra*, from the annotations results we can see *Nvbra* participates in regeneration cluster two (R-2). When exploring the cluster R-2 we see all of the transcripts found in the cluster sorted by membership score. The score reflects how strongly a gene’s expression matches the cluster core. By plotting several high-scoring transcripts we can identify groups of co-expressed genes. For example when we plot NvERTx.4.40781 (best nr hit: XP 015758878.1 forkhead box protein G1-like *Acropora digitifera*), NvERTx.4.57897 (best nr hit: AOP31964.1 dickkopf3-like 1 *Nematostella vectensis*), and NvERTx.4.100808 (best nr hit: AAO27886.2 Brachyury protein *Nematostella vectensis*), we see that they are indeed co-expressed with an initial expression peak between 4 and 16 hours post amputation (hpa), followed by a gradual rise from 36hpa onward (Fig. 4A). By contrast these genes are not co-expressed during embryogenesis and exhibit divergent expression patterns (Fig. 4B) raising the hypothesis that this particular grouping of genes is unique to regeneration. Using this method to find co-expressed genes is an effective way for identifying potential gene-regulatory modules and gene batteries. Importantly, assaying if these genes are co-expressed during regeneration and embryogenesis can shed light on how these gene batteries are used or re-used during these two processes.

**Figure 4:**
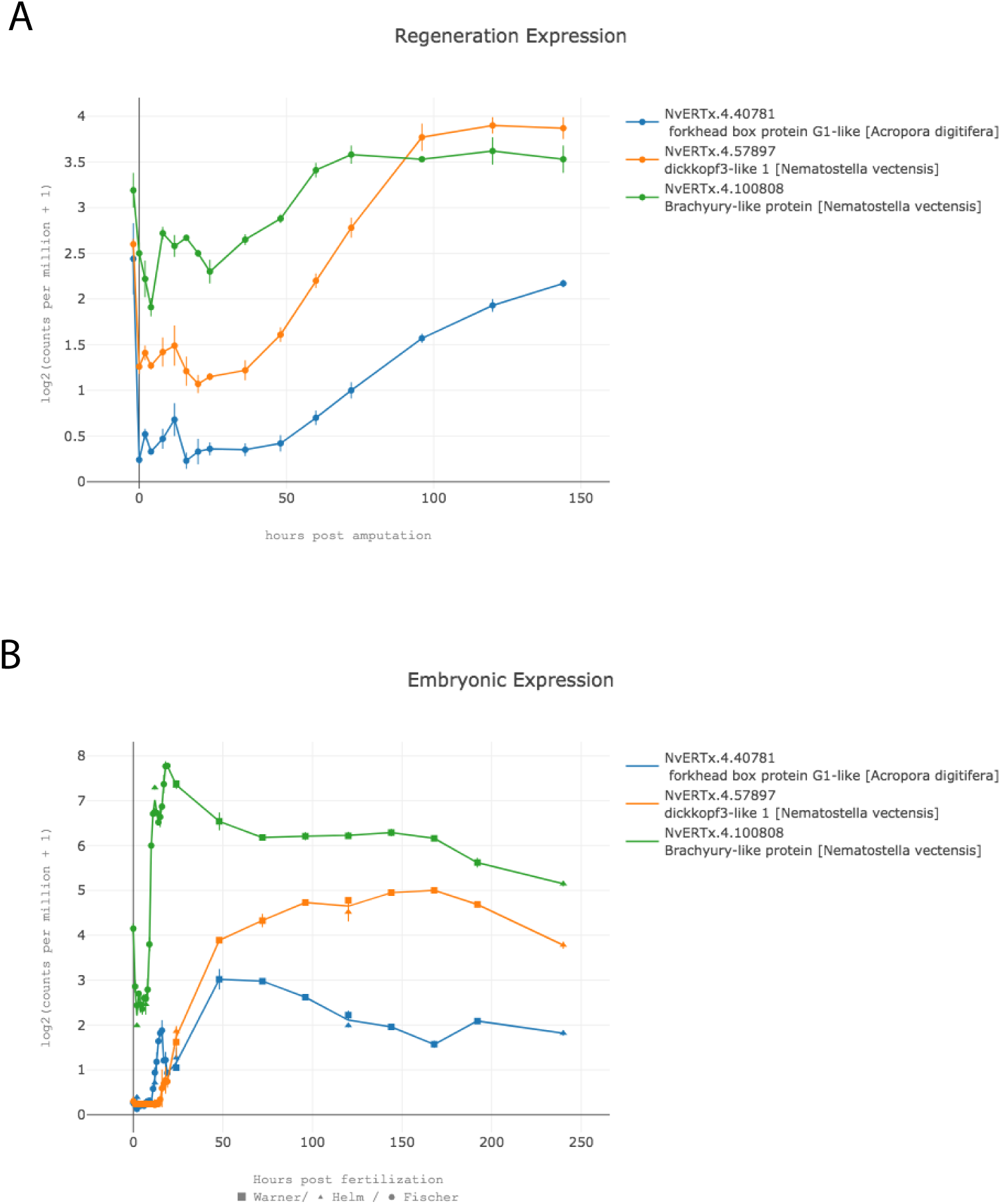
Example output plots from NvERTx comparing multiple gene expression patterns. Three genes from regeneration cluster 2, *Nvbra* (NvERTx.4.100808, yellow), *Nvdickkopf3* (NvERTx.4.57897, red), and a FoxG1-like protein (NvERTx.4.40781, blue) are co-expressed during regeneration (A) but not embryogenesis (B).

**Figure 5:**
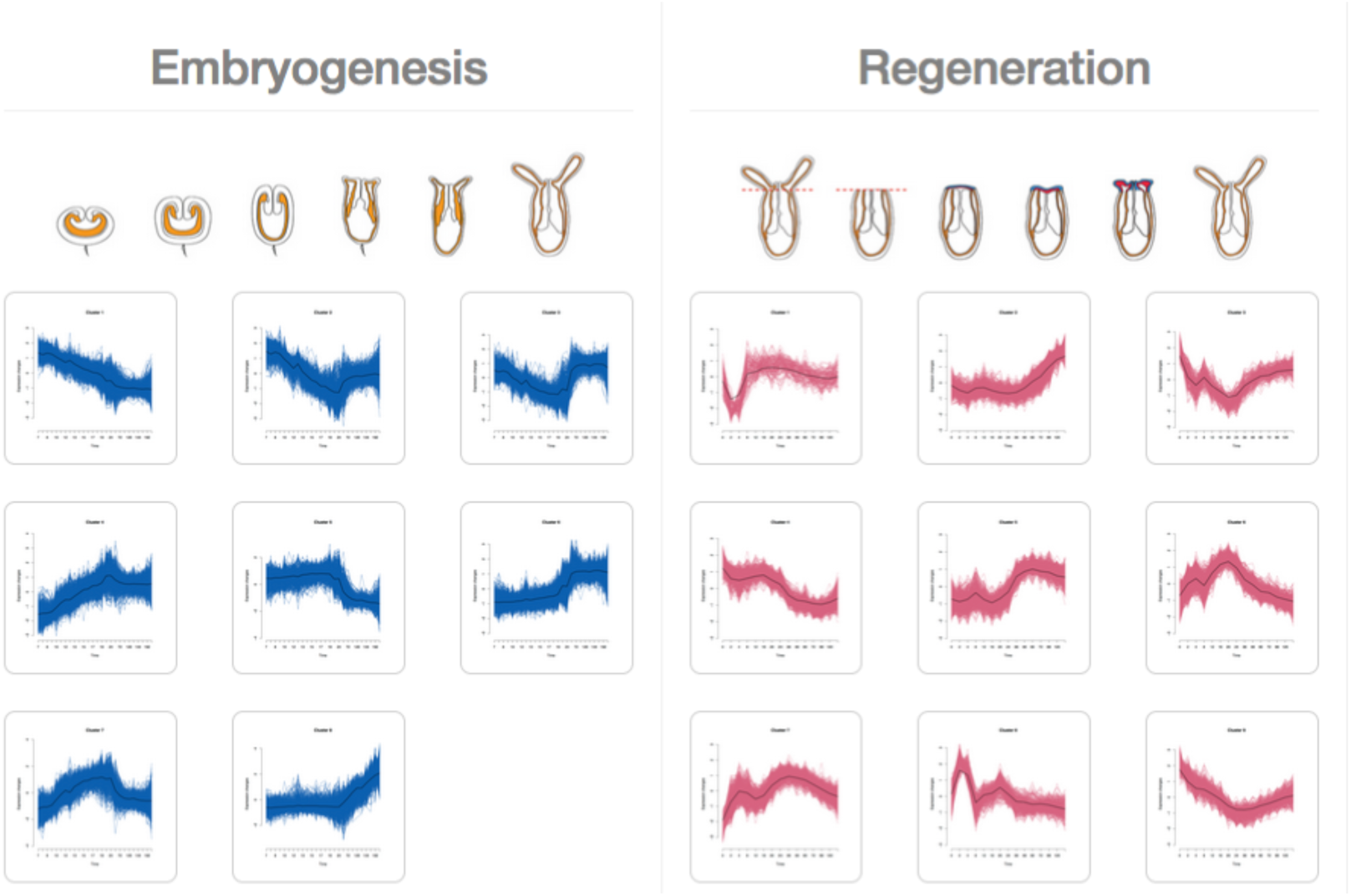
Screenshot from NvERTx Co-Expression Clusters Page. Users can directly explore co-expression clusters to identify groups of genes that share expression patterns during embryogenesis (blue) or regeneration (red).

### Conclusion and future directions

NvERTx provides a platform to efficiently compare gene expression between embryogenesis and regeneration in *Nematostella*. Additionally, the comprehensive transcriptome provides high quality transcript models than can be used to identify gene sequences. Finally, using co-expression clusters, one can explore groups of genes that share similar expression patterns during embryogenesis and regeneration. All of these tools are aimed to inspire and build hypotheses concerning embryogenesis and regeneration for *Nematostella* and *non-Nematostella* researchers alike. Users can use their own groups of co-expressed genes to test for conservation of regeneration gene batteries, or explore the gene expression clusters to identify genes that share expression and examine these in their own models.

As this web application is intended to complement and expand upon existing resources, we provided transcript models that have been annotated using a variety of gene/protein databases (nr, tremble, Nemve1). Sequencing technologies are evolving to achieve longer reads, and assemblers will soon provide more robust transcriptomes and in the future we plan to take advantage of these technologies to improve our transcript models. We also plan to grow the database as future datasets examining embryogenesis and regeneration emerge. Finally, we foresee merging this resource with an existing spatial gene expression database found at http://www.kahikai.org/index.php?content=genes (Ormestad et al., 2011). This will enable identification syn-expression groups, genes which are co-expressed both spatially and temporally (Niehrs and Pollet, 1999), and further facilitate studies comparing differential gene usage during embryogenesis and regeneration.

## Materials and Methods

### Transcriptomic datasets

The sequences that served as input into our *de novo* assembly is comprised of two datasets: One spanning the first 24 hours of embryogenesis originally reported by Tulin et al. 2013 (Tulin et al., 2013) and another spanning the first 144 hours of regeneration generated in house (see library preparation and sequencing). The embryonic dataset was downloaded from the Woods Hole Open Access Server (http://darchive.mblwhoilibrary.org/handle/1912/5613, last accessed June 1, 2017) and includes Illumina HiSeq 100bp paired end sequencing prepared from *Nematostella* embryos at 0, 6, 12, 18, and 24 hours post fertilization. The regeneration dataset includes Illumina NextSeq 75bp paired end sequencing from regenerating *Nematostella* at -1, 0, 2, 4, 8, 12, 16, 20, 24, 36, 48, 60, 72, 96, 120 and 144 hours post amputation (see library preparation and sequencing section for details). The sequence data that was used for transcript quantification is comprised of four datasets. Three datasets spanning embryogenesis: originally reported in Fischer et al. (Fischer et al., 2014) (http://darchive.mblwhoilibrary.org/handle/1912/5981, last accessed June 1, 2017)(Illumina HiSeq 100bp paired end replicates sampled hourly from 0-19 hours post fertilization), the second embryonic dataset originally reported in Helm et al. 2013 (Helm et al., 2013) (NCBI short read archive Project: PRJNA189768)(Illumina HiSeq 50bp single end replicates sampled from 2, 7, 12, 24, 120 and 240), and the third embryonic dataset was generated in house sampled at 24, 48, 72, 96, 120, 144, 168, and 192 hours post fertilization (Illumina MiSeq 75Bp single end replicates). The regenerative data used for quantification is the same used for the transcriptome assembly (Illumina NextSeq 75bp single end triplicates).

### Library preparation and sequencing

Two novel RNAseq datasets, one spanning embryonic, larval and post-metamorphic development and the other regeneration were generated for this study:

For the embryonic dataset, approximately 250 embryos per time point were cultured in 1/3 strength artificial seawater at 18°C. At each time point, 24, 48, 72, 96, 120, 144, 168, and 192 hours post fertilization, the embryos were transferred to 500ml of Tri-Reagent and homogenized for 15 seconds with a pistle. The resulting lysate was snap frozen in liquid nitrogen and stored at -80°C. This was repeated to obtain duplicates for each time point. After all the samples were collected, the RNA lysate was extracted using two phenol-chloroform extractions and precipitated with isopropanol. For the full extraction protocol, see: (Layden et al., 2013). The resulting nucleic acids were treated with the TURBO DNA-free kit from Invitrogen (AM1907) for 10 minutes at 37°C. The resulting RNA was quantified with a Qubit spectrometer and RNA integrity was checked on an Agilent Bioanalyzer 2100. 100ng of RNA was used as input to the Illumina Truseq stranded mRNA library Prep for Neoprep kit and the libraries were prepared using the automate Illumina Neoprep system. 75Bp single end sequencing was carried out on the NextSeq500 sequencer

For the regenerative dataset, approximately 350 six-week-old Nematostella juveniles per time point were amputated below the pharynx. At each time point, -1, 0, 2, 4, 8, 12, 16, 20, 24, 36, 48, 60, 72, 96, 120, and 144 hours post amputation, the juveniles were transferred to 500ml of Tri-Reagent and homogenized for 15 seconds with a pistle. The resulting lysate was snap frozen in liquid nitrogen and stored at -80°C. This was repeated for each of the three replicates. After all the samples were collected the RNA was extracted as described above for the embryonic samples. Samples were stored in GenTegra-RNA stabilization reagent (prudct ID: GTR5100-S) and shipped to the NextGen Sequencing Core at the University of Southern California for library preparation and sequencing. The samples were prepared with KAPA stranded RNA kit (KR0960). Two replicates were sequenced as single end 75Bp and one replicate as paired end 75Bp on an Illumina NextSeq500 sequencer.

### RNAseq quality control

All reads from each dataset were processed equivalently. Reads were first quality filtered to remove low quality reads and adapter trimmed using timmomatic (Bolger et al., 2014) and cutadapt (MARTIN, 2011) respectively.

### Trinity de novo assembly

For the *de novo* assembly, paired end reads from regeneration (-1, 2, 4, 8, 12, 16, 20, 24, 36, 48, 60, 72, 96, 120 and 144 hpa) and embryonic dataset (0, 6, 12, 18, and 24 hpf, (Tulin et al., 2013)) were filtered of ribosomal sequences by aligning to *Nematostella* mitochondrial and ribosomal sequences using Bowtie2 (Langmead et al., 2009) and retaining the unmapped reads. These surviving reads were input into Trinity (v2.4.0) for assembly (Haas et al., 2013). To annotate the assembly the ORFs of the open reading frame were found using OrfPredictor (Min et al., 2005) and the resulting peptide sequences were compared to the NCBI nr database using PLAST (Nguyen and Lavenier, 2009) with an evalue cutoff of 5e-5. The transcriptome was also compared to the Uniprot databases, swissport and trembl, using BLASTx and PLASTp respectively with an evalue cutoff of 5e-5.

### Quantification

To quantify the RNAseq data, single end reads for each dataset, regeneration and the three embryonic datasets Fischer, Helm and this study, were aligned to the Trinity assembly using Bowtie2 (Langmead et al., 2009). Read counts were quantified using RSEM (Li and Dewey, 2011). To account for the many isoforms per gene reported by Trinity, transcripts were compared to the Nemve1 filtered gene models using BLASTn and counts for transcripts with the same Nemve1 hit were combined. Transcripts with low read counts, those that did not have >5 counts in at least 25% of the samples, were excluded. Each dataset was then normalized separately using the R package edgeR and the counts per million (cpm) mapped reads were calculated. To correct for batch effects of the embryonic datasets after normalization, the R function ComBat from the SVA packages was used on log2(cpm+1) transformed data using timepoint as a categorical covariate (Leek et al., 2012).

### Fuzzy c-means clustering and GO term enrichment

For each Nemve1 gene model, transcriptional expression profiles were clustered using the R package mFuzz (Kumar and E Futschik, 2007). The cluster number was set to 9 for the regeneration data and 8 for the embryonic datasets as these numbers represented the point at which the centroid distance between clusters did not significantly decrease when new clusters were added (inflection point). For each cluster, GO term enrichment was calculated using a Fisher’s exact test and the R package topGO on the GO terms identified from comparing the transcriptome to the Uniprot database (all identified GO terms were used as a background model). The resulting GO term list was reduced and plotted using a modified R script based on REVIGO (Supek et al., 2011).

### NvERTx website

The website was constructed using the Django python framework (https://www.djangoproject.com/). Plots are generated using the Plotly javascript library (https://plot.ly/javascript/). The source code for the website can be found at https://github.com/IRCAN/NvER_plotter_django. Datasets from the database can be found at http://ircan.unice.fr/ER/ERplotter/about.

## Acknowledgements

- We thank Valerie Carlin for animal care.
- We thank Christian Bauduin for library preparation and the Ircan genomic core facility for mRNAseq libraries preparation and sequencing.

## Author contributions

ER and JFW designed the study; JFW, HJ, KN and ARA collected samples; JFW analyzed data; JFW and VG coded the website; JFW and ER wrote the manuscript. All authors read and approved the final manuscript.

## Data availability

Original datasets generated for this study are found on the NCBI SRA BioProject accessions: PRJNA418421 and PRJNA419631.

